## Supplementary Information File for "Structure of the human CTF18–RFC clamp loader bound to PCNA"

Briola G. R. *et al*

Includes:

Supplementary Figures 1-15

Supplementary Table 1

Supplementary Video 1

References

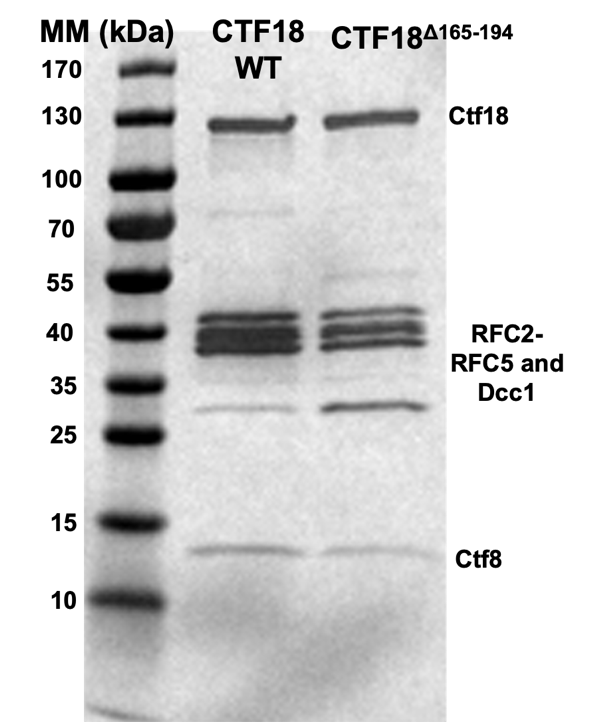

**S1 Fig.** *SDS-PAGE gel (4-20%) of the purified human clamp loader CTF18-RFC WT and CTF18^Δ165-194^*-RFC. Bands corresponding to the different subunits are labeled. The molecular weight standard is shown.

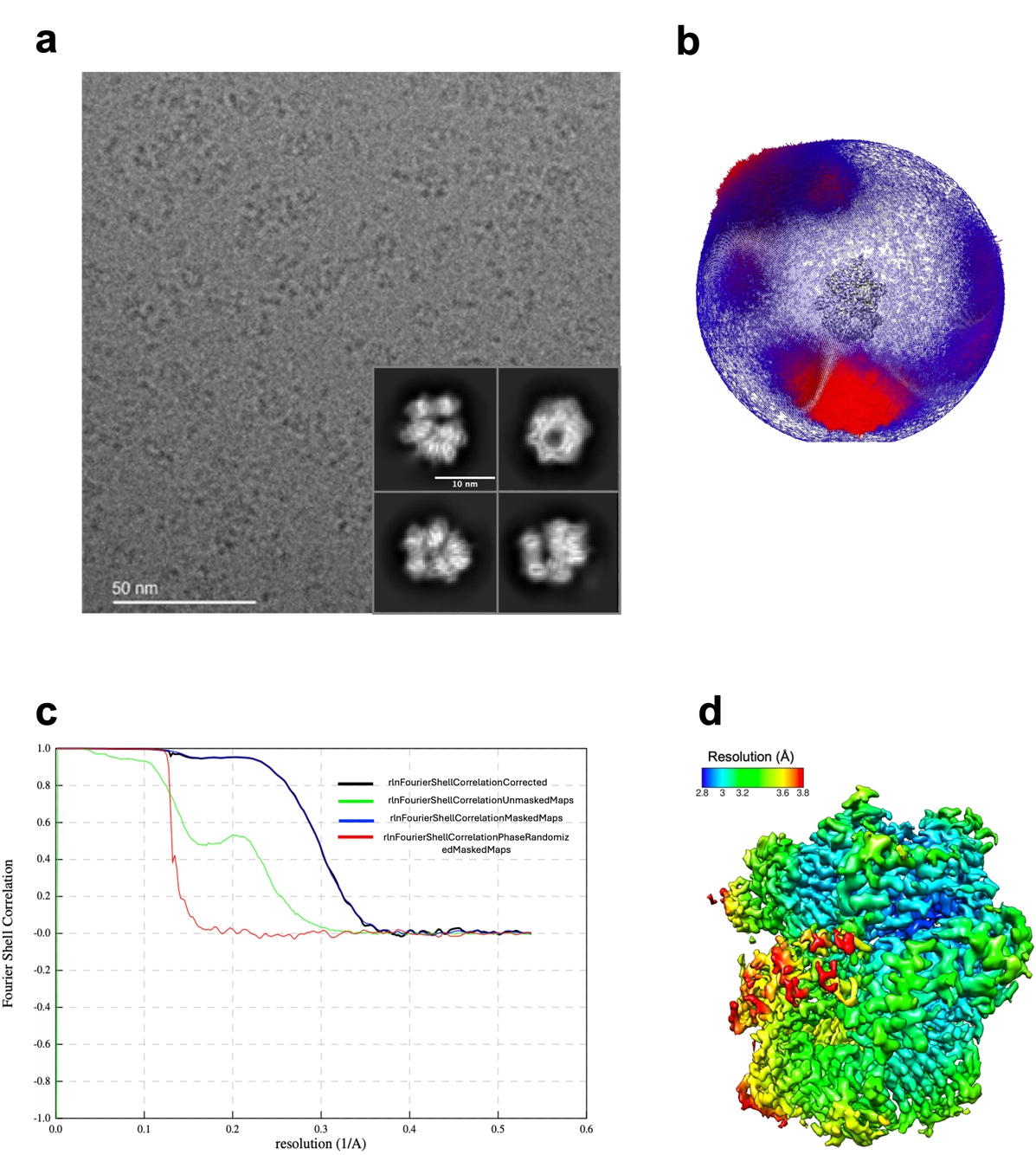

**S2 Fig.** *Cryo-EM of the CTF18−PCNA complex in the presence of ATP.* **a)** Representative electron micrograph acquired on a Falcon 4i electron detector in counting mode, and representative 2D class averages. **b)** Angular distributions of projections. **c)** Gold-standard Fourier shell correlation for the reconstruction of the full complex after postprocessing, and resolution estimation using the 0.143 criterion. **d)** Cryo-EM map of the complex, color-coded by local resolution.

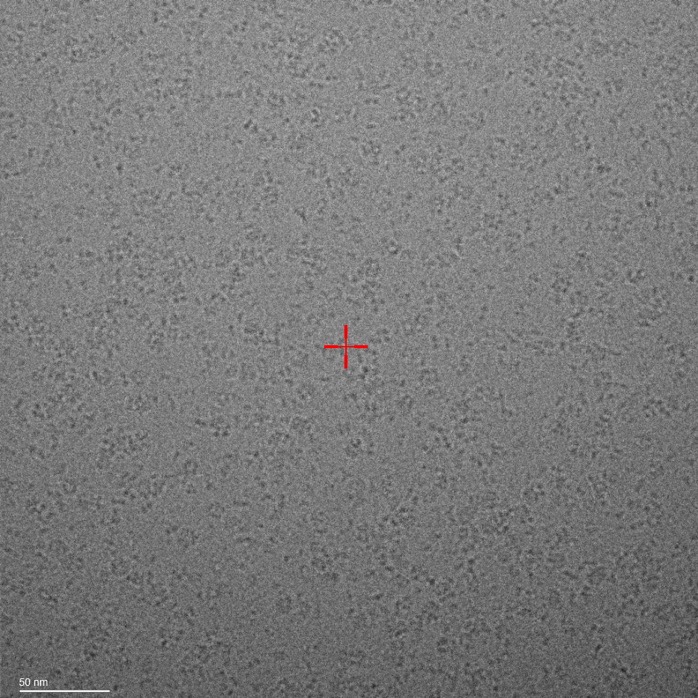

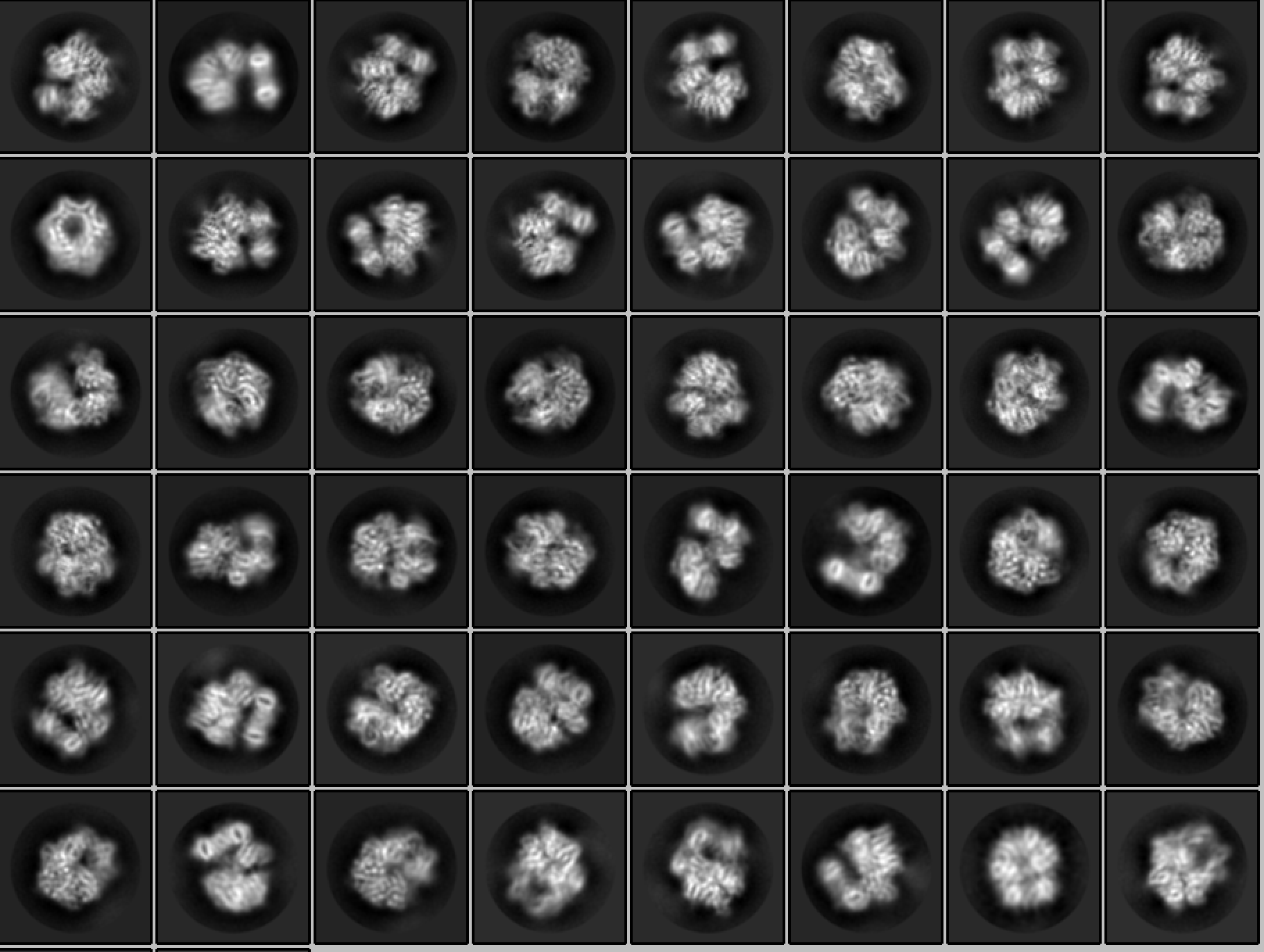

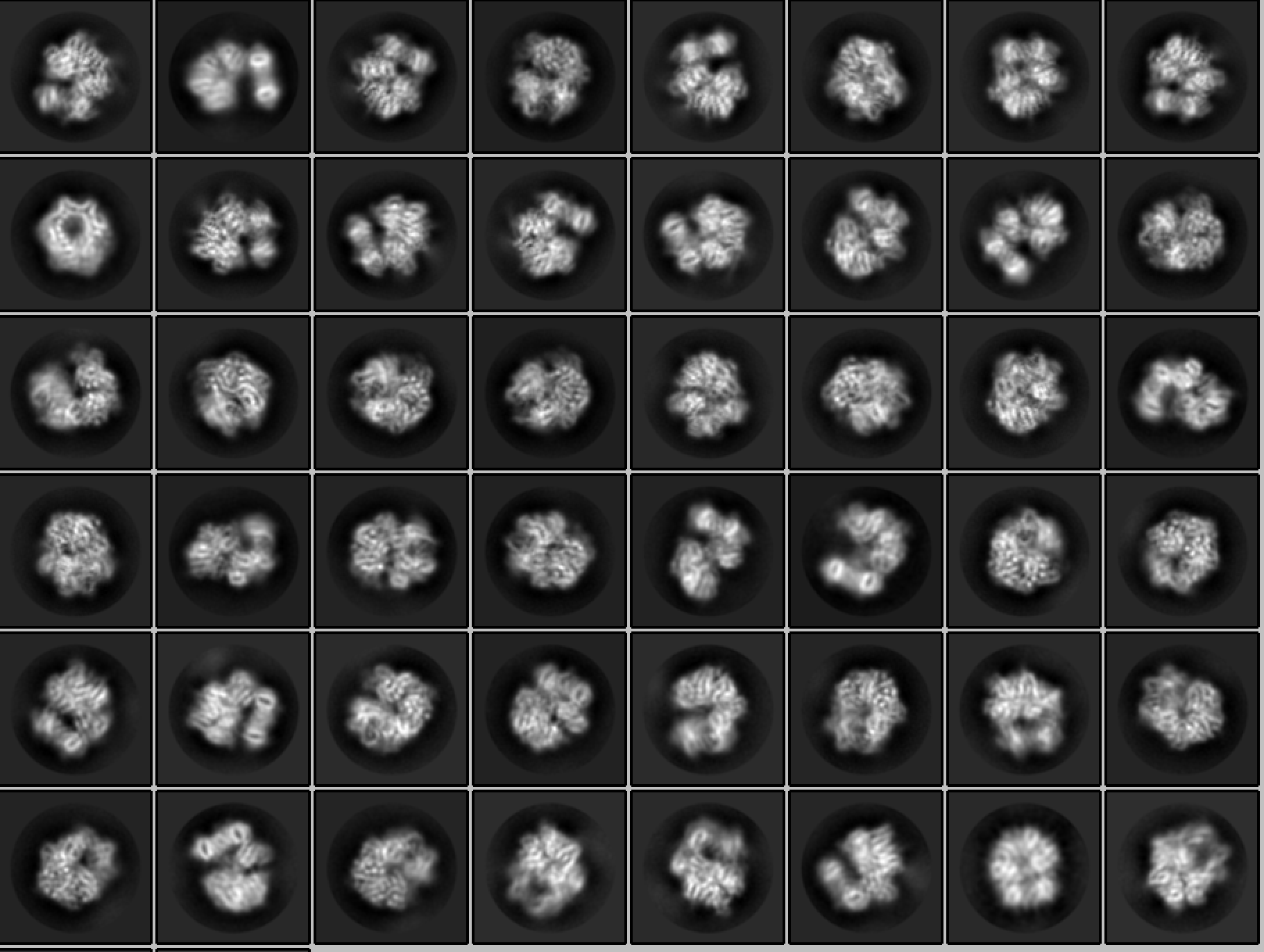

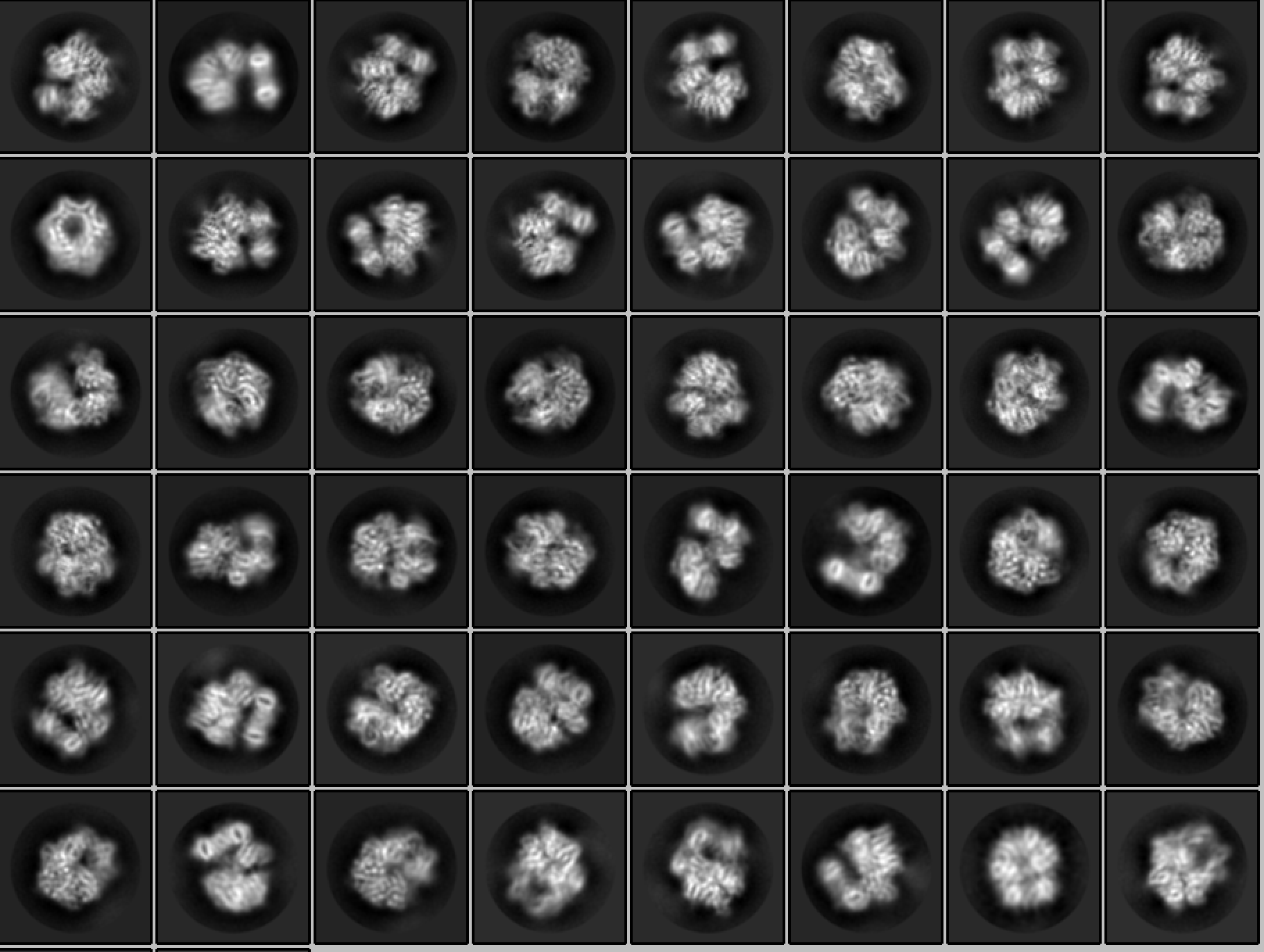

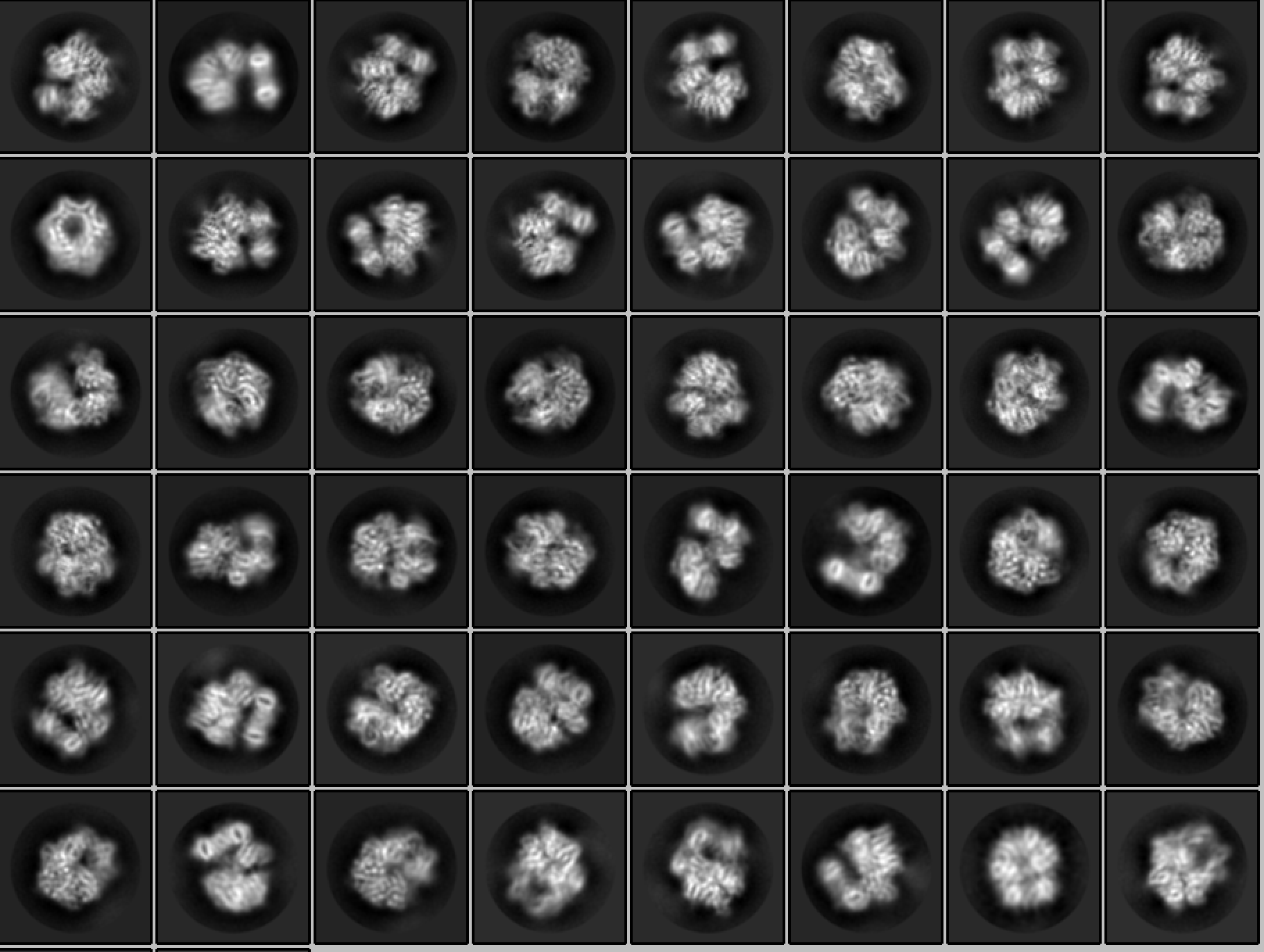

**220 pix**

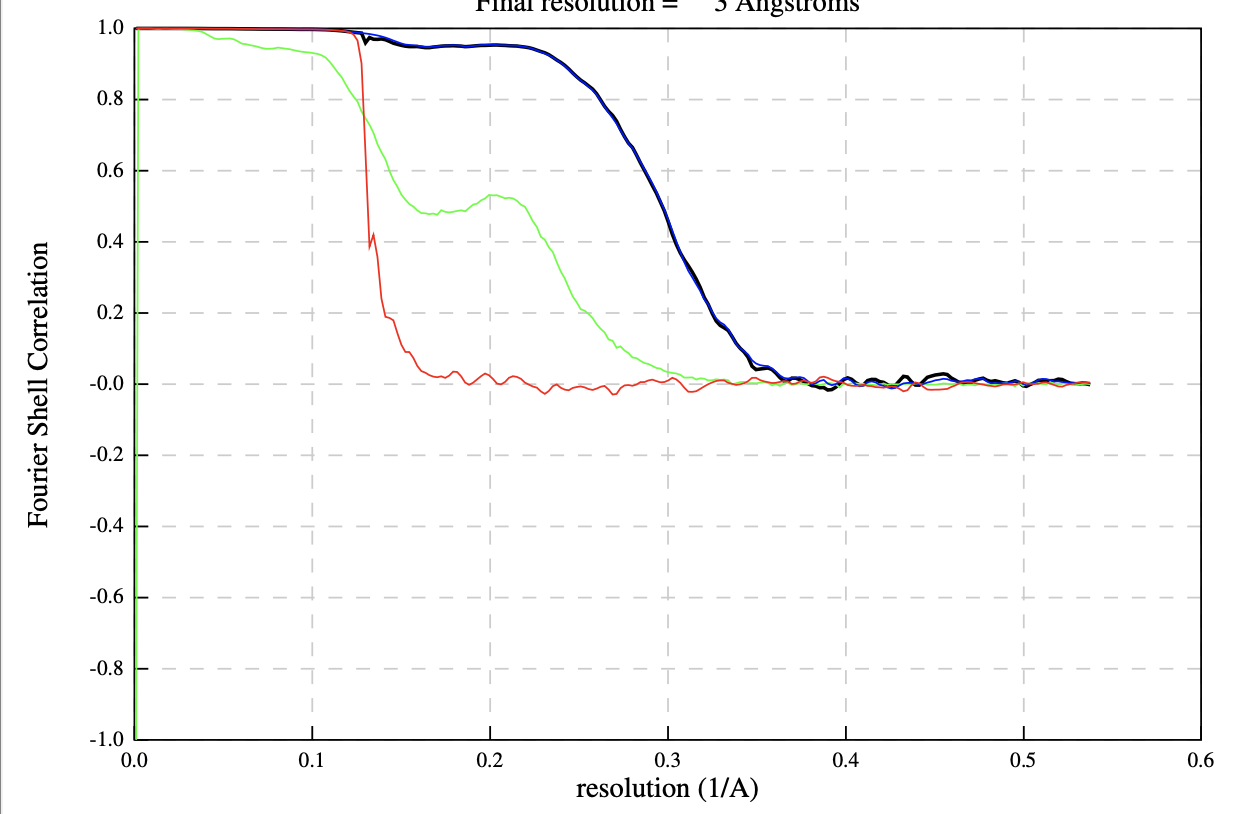

rlnFourierSchellCorrelationCorrected

rlnFourierSchellCorrelationUnmaskedMaps

rlnFourierSchellCorrelationMaskedMaps

rlnFourierSchellCorrelationPhaseRandomizedMaskedMaps

**a**

**b**

**c**

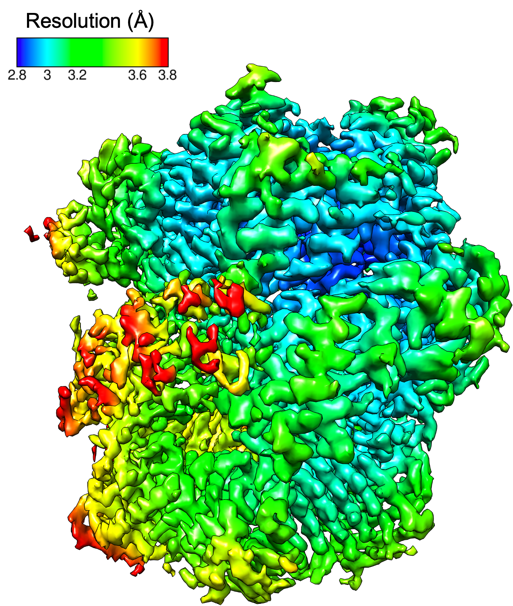

**d**

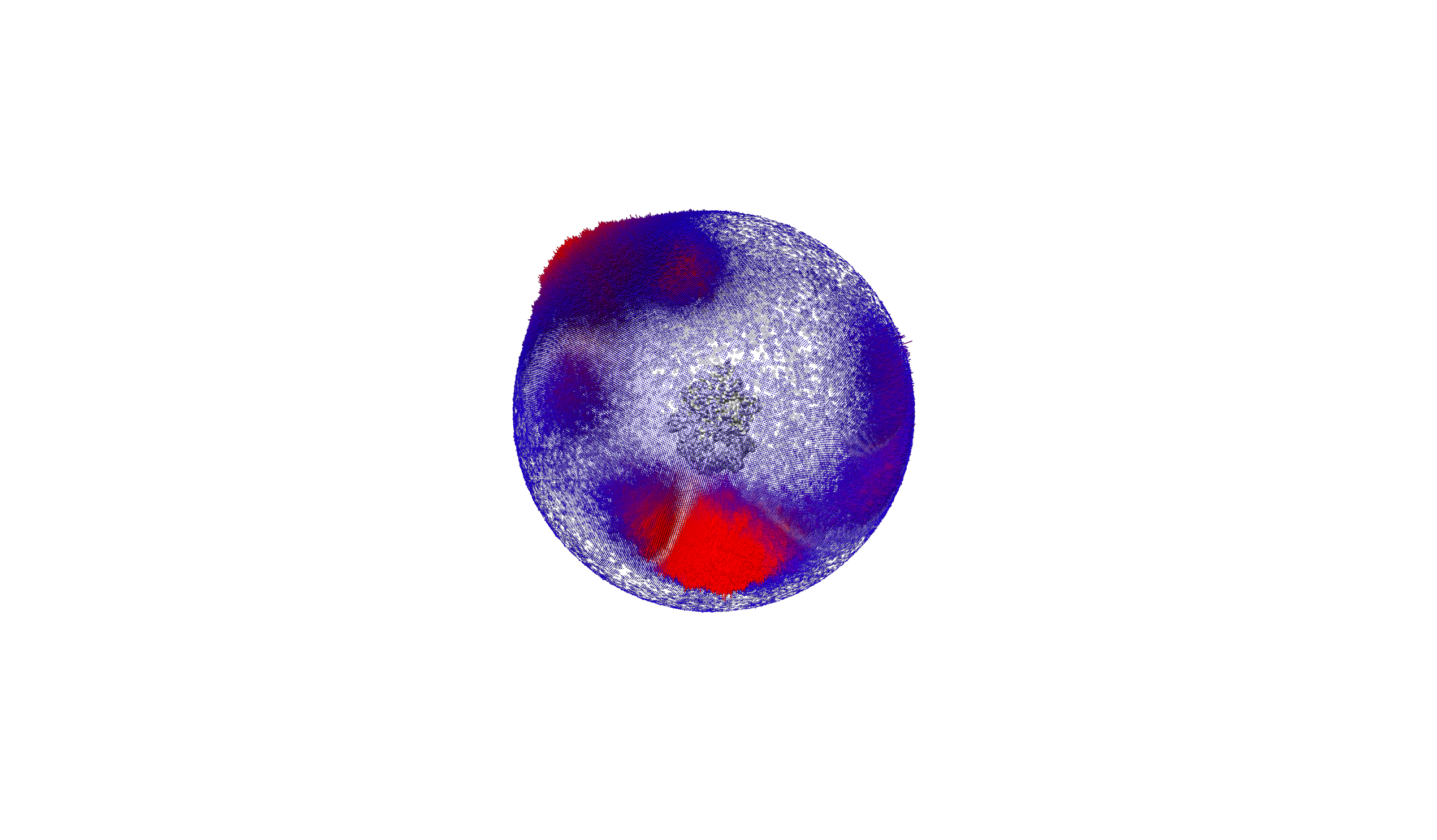

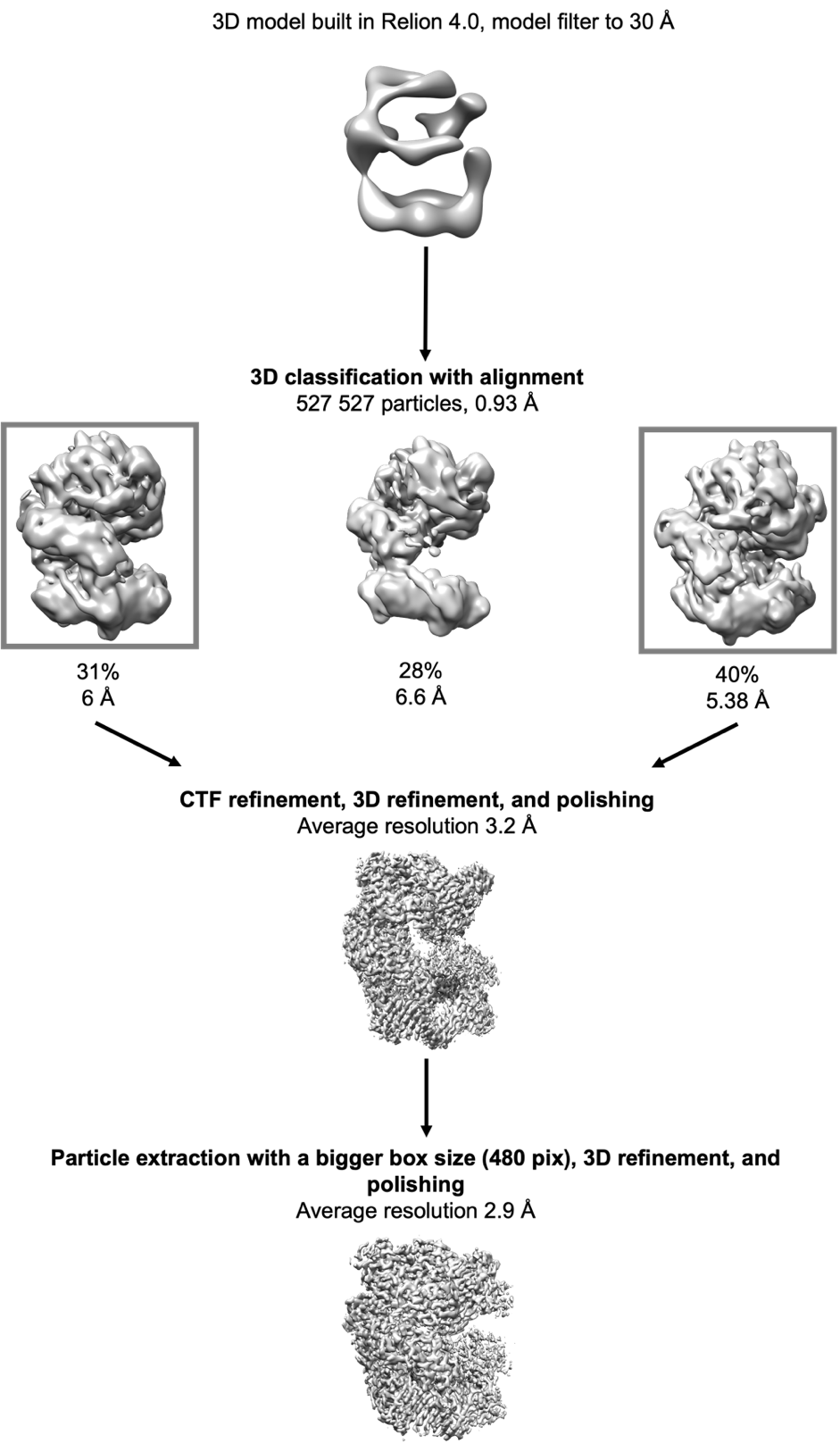

**S3 Fig.** Workflow of cryo-EM image processing and 3D reconstruction of CTF18-RFC-PCNA complex in the presence of 0.5 mM ATP and without Mg^2+^ **(Dataset 1)**. Relion 4.0 was used for image processing and 3D reconstruction.

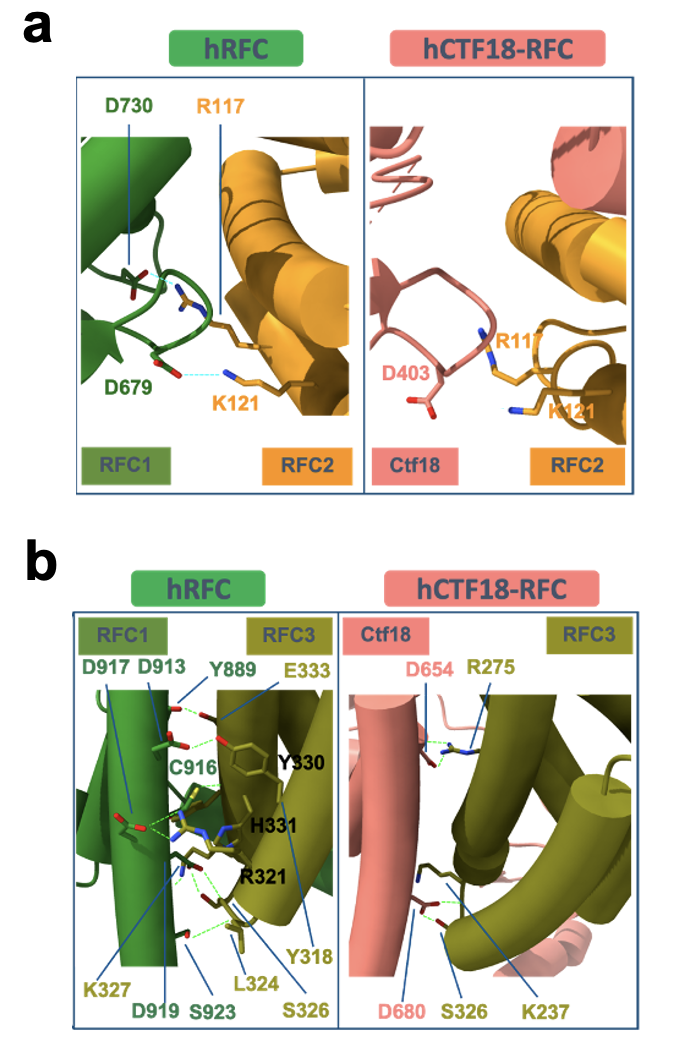

**S4 Fig:** *Interfaces between RFC1 or Ctf18 and RFC2, and RFC1 or Ctf18 and RFC3.* **(a)** Comparison of the AAA+ module interfaces. Key interactions observed with RFC1 are lost with Ctf18 **(b)** Comparison of the collar module interface. Far fewer interactions are observed with Ctf18 compared to RFC1. Salt bridges are shown in turquoise and h-bonds in green.

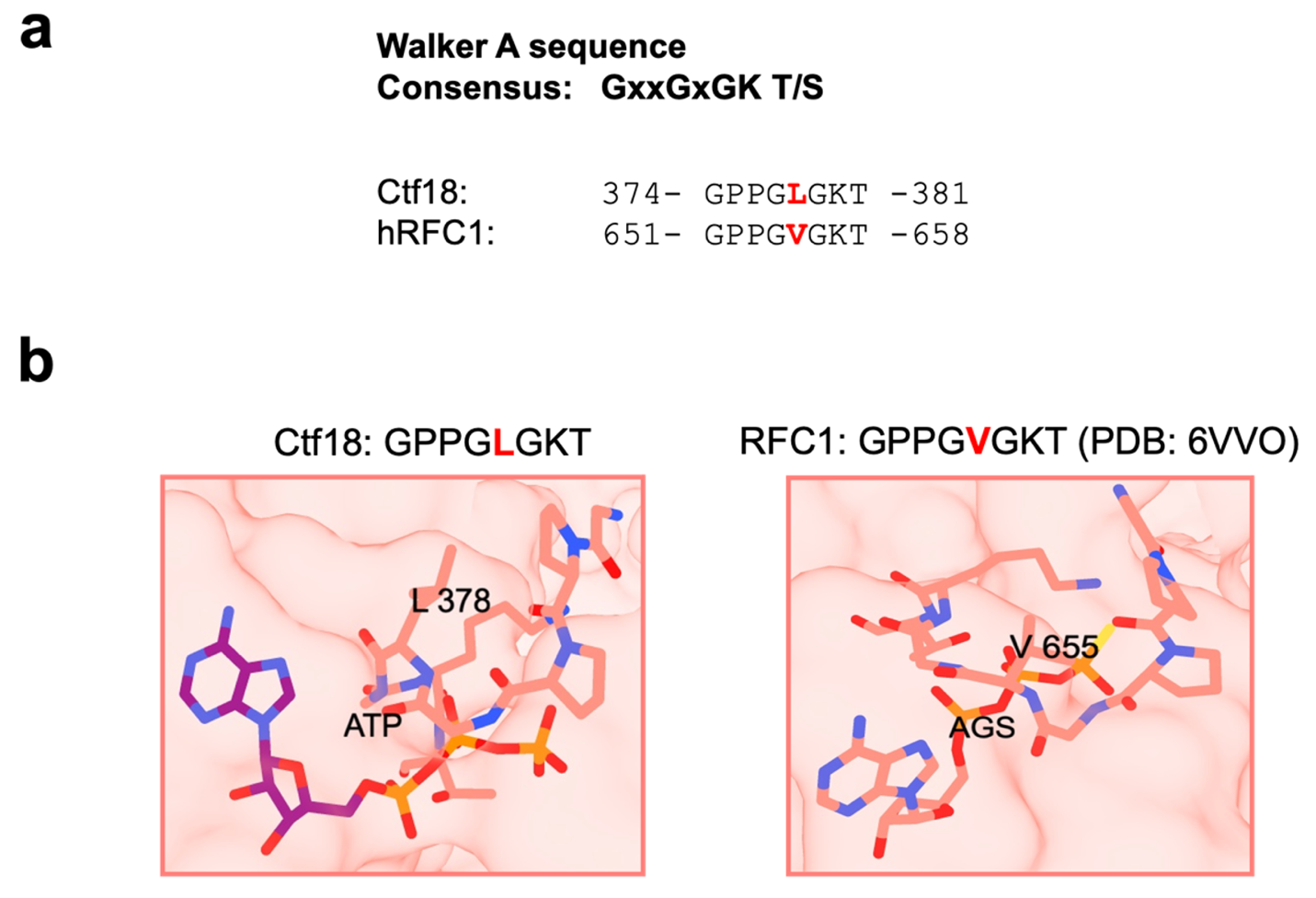

**S5 Fig.** *Walker A sequence and structure.* **a)** Walker A sequence comparison between CTF18 and hRFC (PDB: 6VVO). **b)** The upper panels illustrate the conformation of the Walker A motifs in Ctf18 and human RFC1 (PDB: 6VVO).

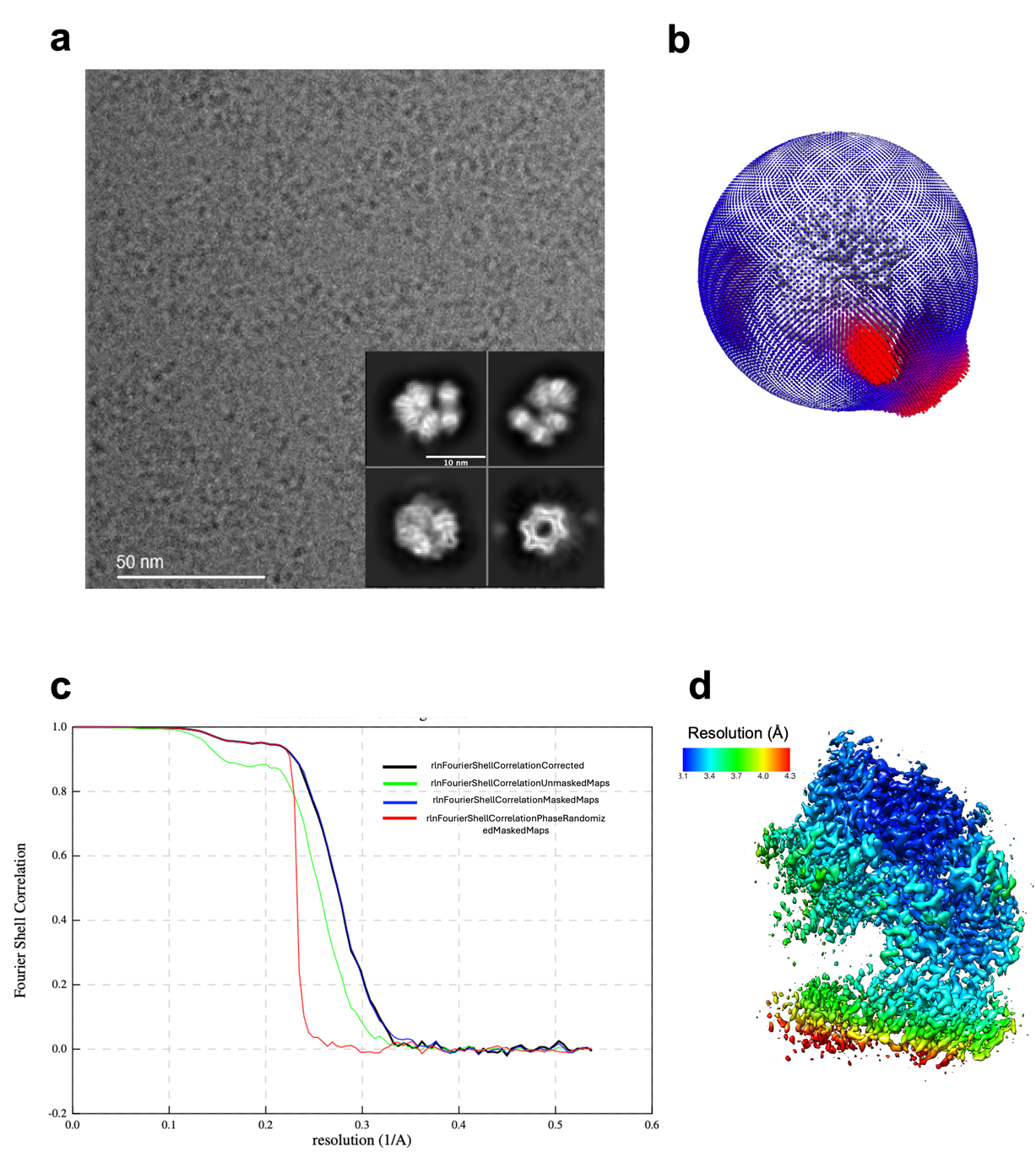

**S6 Fig.** *Cryo-EM of the CTF18−PCNA complex in the presence of ATP and Mg^2+^.* **a)** Representative electron micrograph acquired on a Falcon 4i electron detector in counting mode, and representative 2D class averages. **b)** Angular distributions of projections. **c)** Gold-standard Fourier shell correlation for the reconstruction of the full complex after focused refinement, and resolution estimation using the 0.143 criterion. **d)** Cryo-EM map of the complex, color-coded by local resolution.

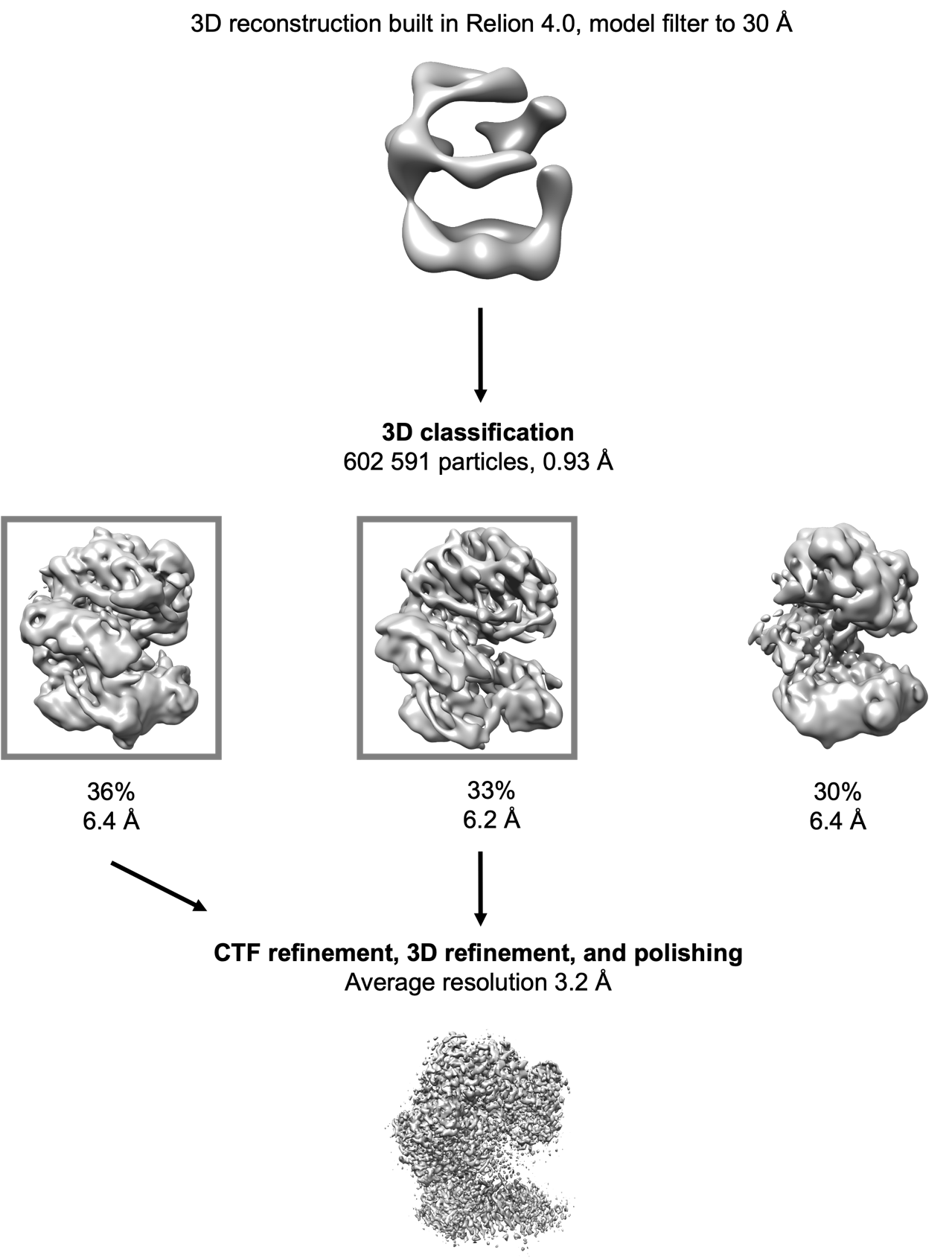

**S7 Fig.** Workflow of cryo-EM image processing and 3D reconstruction of CTF18-RFC-PCNA complex in the presence of 0.5 mM ATP and 5 mM Mg^2+^ **(Dataset 2)**. Relion 4.0 was used for image processing and 3D reconstruction.

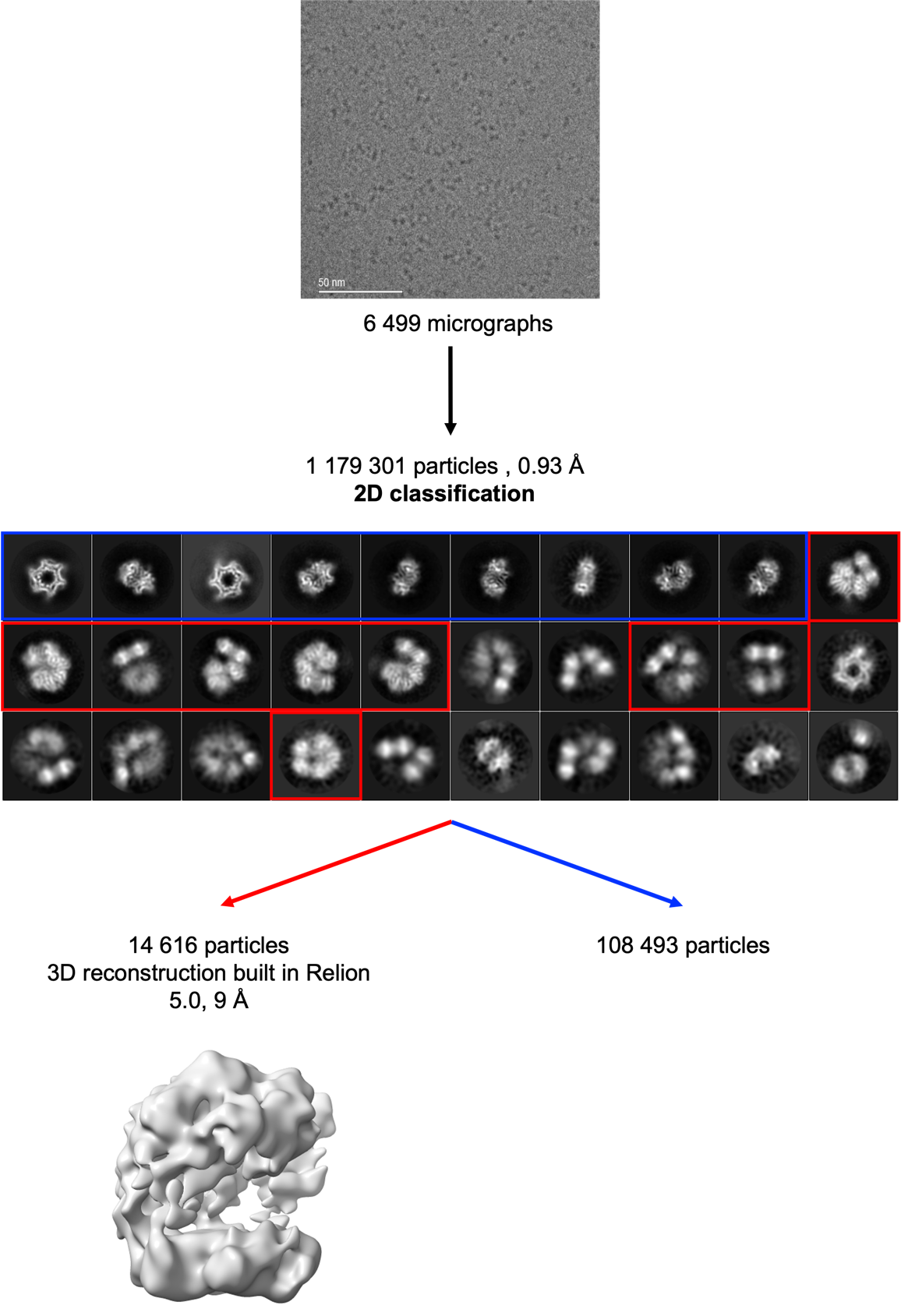

**S8 Fig.** Workflow of cryo-EM image processing and 3D reconstruction of *CTF18-RFC ^Δ165-194^*-PCNA complex in the presence of 0.5 mM ATP and without Mg^2+^ **(Dataset 3)**. Relion 5.0 was used for image processing and 3D reconstruction.

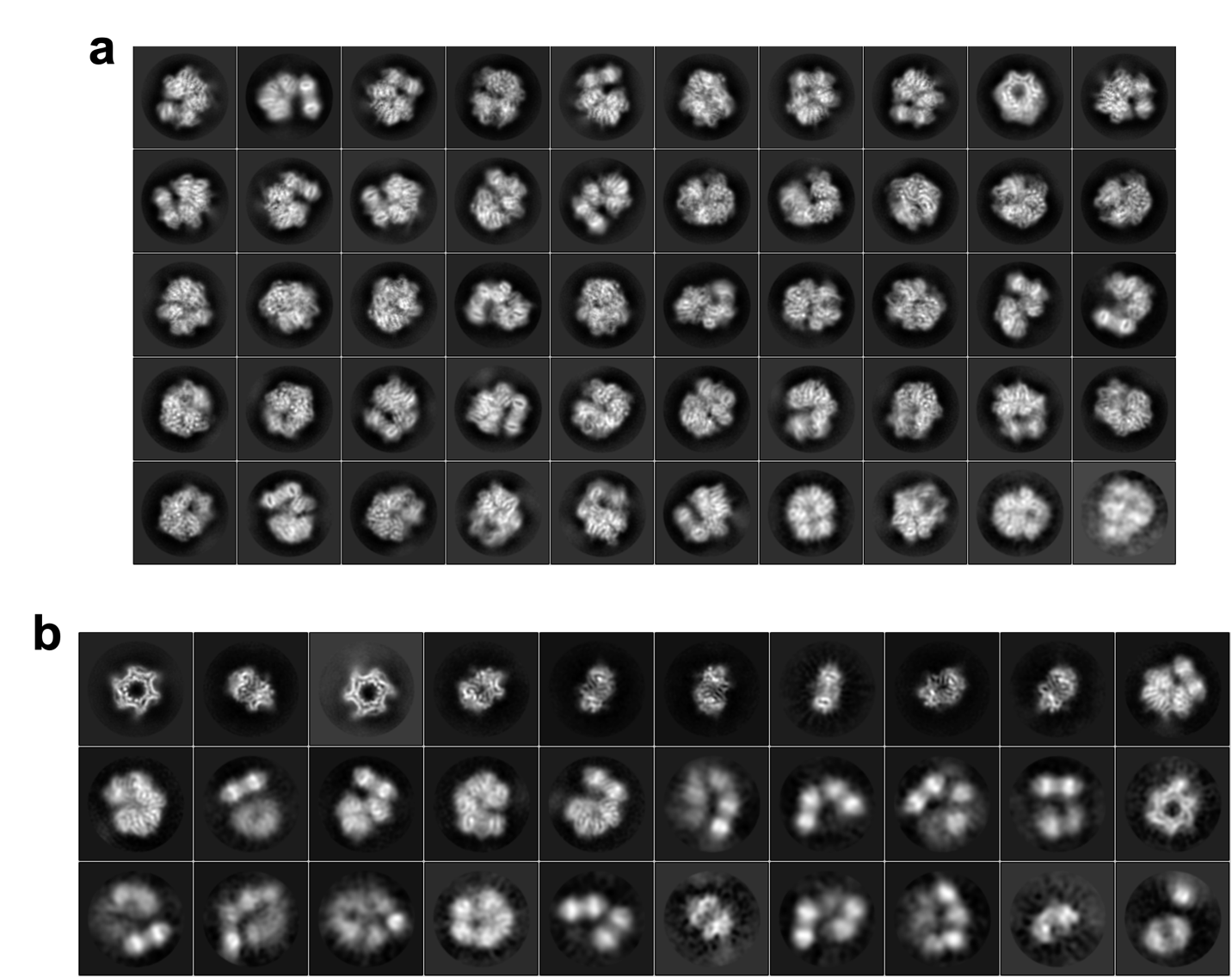

**S9 Fig.** *2D classification of CTF18 WT and CTF18^Δ165-194^-RFC.* **a)** The 2D classification of CTF18-RFC WT reveals the formation of the CTF18-RFC−PCNA complex with clear structural integrity (515,195 particles in all the showed 2D classes) **b)** The 2D classification of CTF18 *^Δ165-194^*-RFC mutant highlights the impact of β-hairpin deletion on complex stability. The presence of a large majority of particles of PCNA alone (108,439 particles) indicates that the absence of the β-hairpin contributes to the increased instability of the complex. Only the 2D classes squared in red represent the CTF18 *^Δ165-194^*-RFC−PCNA complex (14,616 particles).

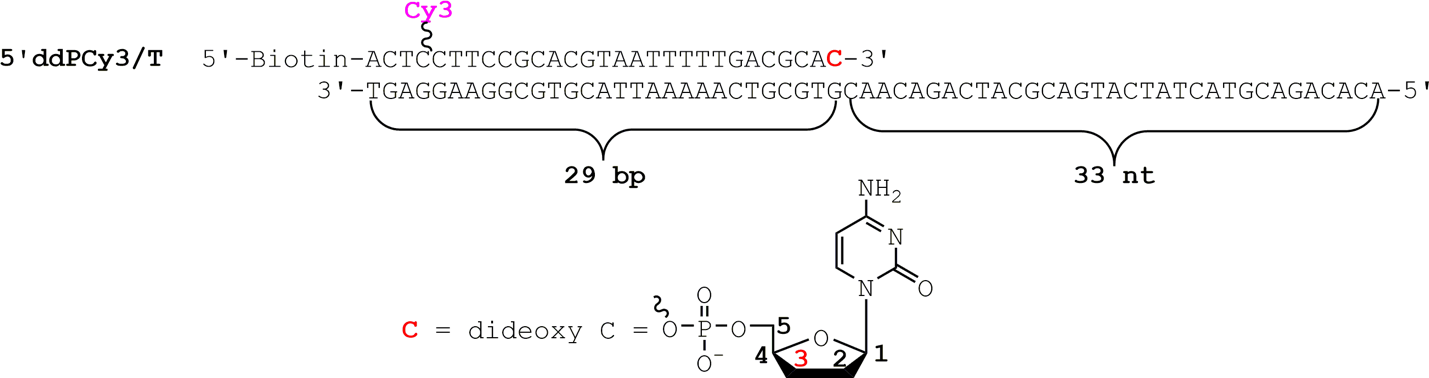
**S10 Fig*.*** *P/T DNA substrate utilized in the FRET studies*. The sequences and lengths of the double- and single-stranded DNA regions are indicated. The size of the double-stranded DNA (dsDNA) region (29 bp) is in agreement with the requirements for assembly of a PCNA ring onto DNA by RFC (1, 2, 3). The single-stranded DNA (ssDNA) region accommodates 1 RPA heterotrimer (4, 5, 6). RPA prevents loaded PCNA from sliding off the ssDNA end of the substrate (2). When pre-bound to neutravidin, the biotin attached to the 5¢-end of the primer strand of the substrate prevents loaded PCNA from sliding off the dsDNA end. The primer strand is terminated at the 3¢ end with a dideoxy C nucleotide (shown) and, hence, cannot be extended by a DNA polymerase.

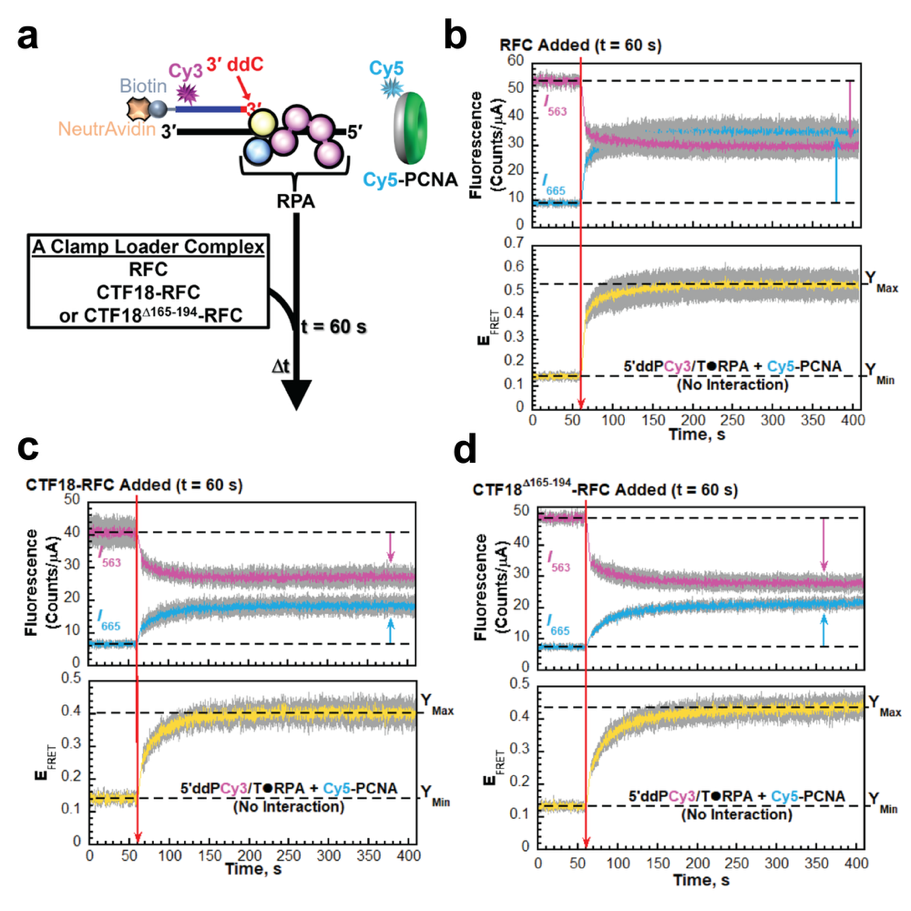

**S11 Fig.** *FRET assays to monitor loading of PCNA onto P/T junctions by human clamp loader complexes*. **a)** Schematic representation of the FRET pair and experiment to monitor loading of Cy5-PCNA onto 5′ddPCy3/T DNA substrates engaged by RPA. The front and back faces of PCNA are displayed in green and grey, respectively. When loaded onto a 5′ddPCy3/T DNA substrate, the Cy5 label on the back face of PCNA is oriented towards the Cy3 label near the blunt duplex end of the 5′ddPCy3/T DNA substrate, yielding a FRET signal **b-d)** Data. Each trace is the mean of at least three independent traces with the S.E.M. shown in grey. The time trajectories of I_563_ (magenta) and I_665_ (cyan) are displayed in the top panels and the corresponding E_FRET_ (mustard) is displayed in the bottom panels. The time at which a clamp loader complex is added (t = 60 s) is indicated. by a red arrow. Changes in I_563_ and I_665_ are indicated in the top panel by magenta and cyan arrows, respectively. The I_563_, I_665_, and the corresponding E_FRET_ values observed prior to the addition of clamp loader complexes represents the complete absence of interactions between 5¢ddPCy3/T·RPA complexes and Cy5-PCNA. These values are each fit to flat line that is extrapolated to the axis limits where the Y-intercept of the fit for the E_FRET_ is equivalent to Y_Min_ (indicated). The E_FRET_ values observed over the last 60 s of the plateaus for the observed E_FRET_ increases are each fit to flat line that is extrapolated. to the axis limits where the Y-intercepts are equivalent to Y_Max_ (indicated). Data for loading of PCNA by RFC, CTF18, and CTF18^Δ165-194^-RFC are displayed in panels **b**, **c**, and **d**, respectively.

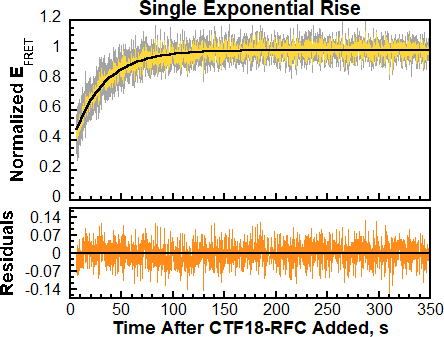

**S12 Fig**. The minimal kinetic model for the increase observed in the normalized E_FRET_ trace for CTF18 is a single exponential rise. The normalized E_FRET_ trace is displayed in the top panel (in yellow) and fit to a single exponential rise. The trace is the mean of at least three independent traces with the S.E.M. shown in grey. The residuals from the corresponding standard curve fitting of normalized E_FRET_ trace are displayed in the bottom panel (in orange).

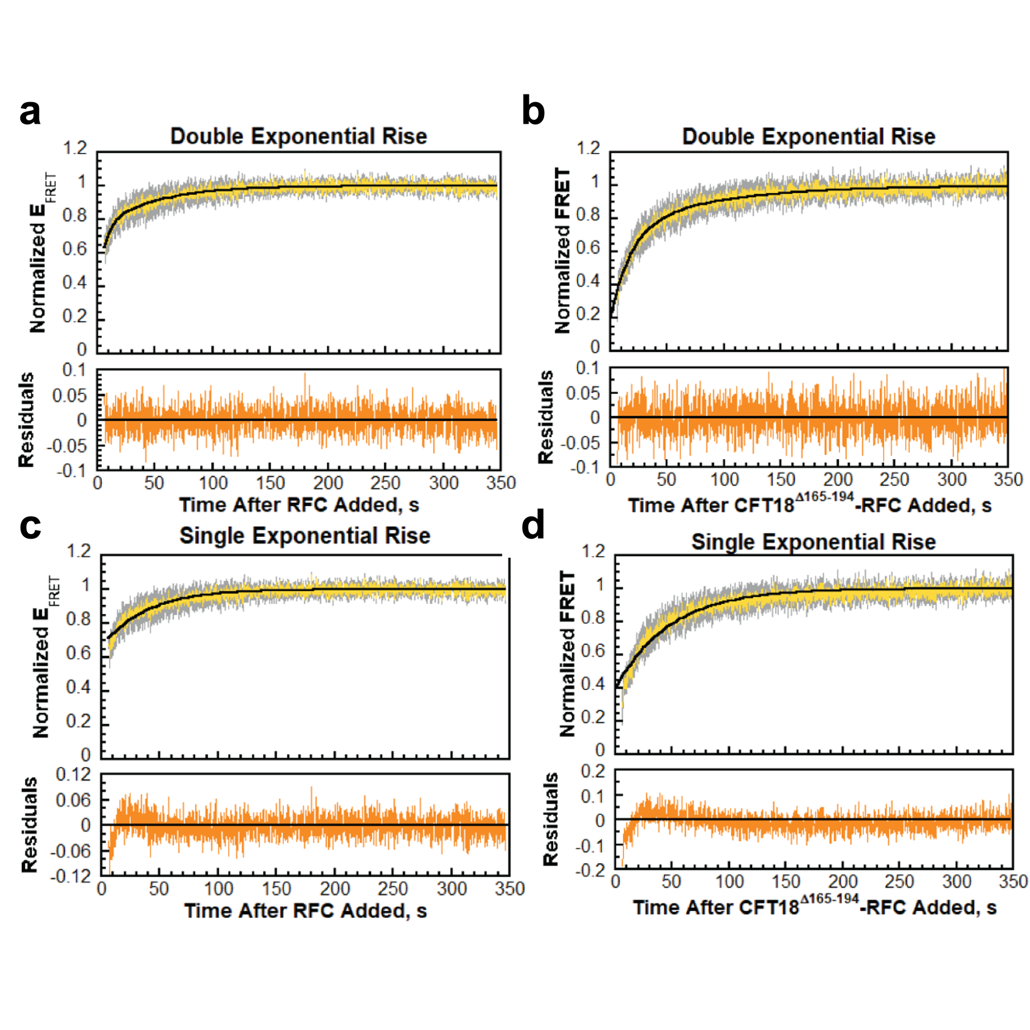

**S13 Fig.** The minimal kinetic models for the increase observed in the normalized E_FRET_ traces for RFC and CTF18^Δ165-194^-RFC are double exponential rises. The normalized E_FRET_ traces are displayed in the top panels (in yellow) and fit to a kinetic model (indicated). Each trace is the mean of at least three independent traces with the S.E.M. shown in grey. The respective residuals from the standard curve fittings of the corresponding normalized E_FRET_ traces are displayed in the bottom panels (in orange). Standard curve fittings of the normalized E_FRET_ traces for RFC and CTF18^Δ165-194^-RFC to double exponential rises are shown in **a** and **b**, respectively. Standard curve fittings of the normalized E_FRET_ traces for RFC and CTF18^Δ165-194^-RFC to single exponential rises are shown in **c** and **d**, respectively.

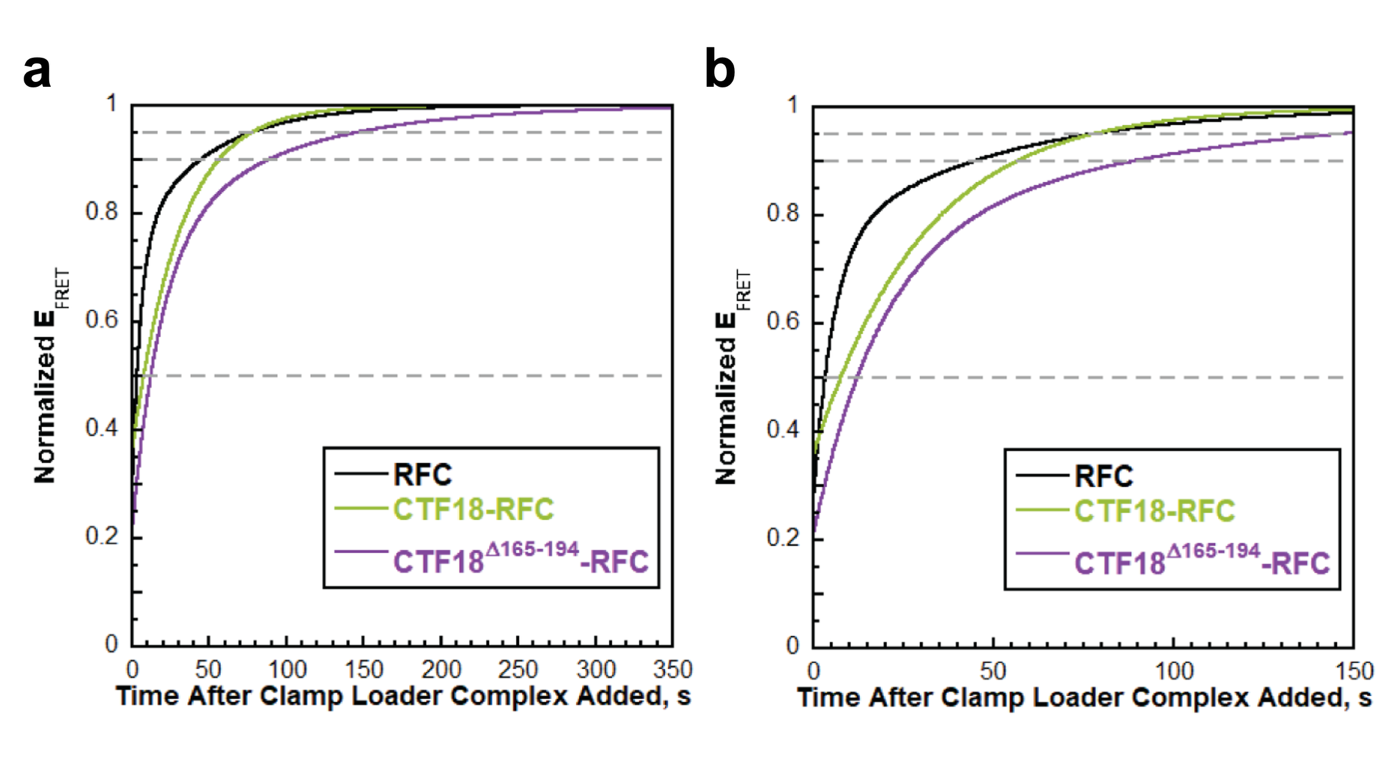

**S14 Fig**. Direct comparisons of the minimal kinetic models for the increase observed in the normalized E_FRET_ traces for RFC (Black), CTF18-RFC (Green), and CTF18^Δ165-194^-RFC (Purple). For perspective, dashed, flat grey lines at 0.5, 0.9 and 0.95 The time courses to 350s and 150 s are displayed in **a** and **b**, respectively.

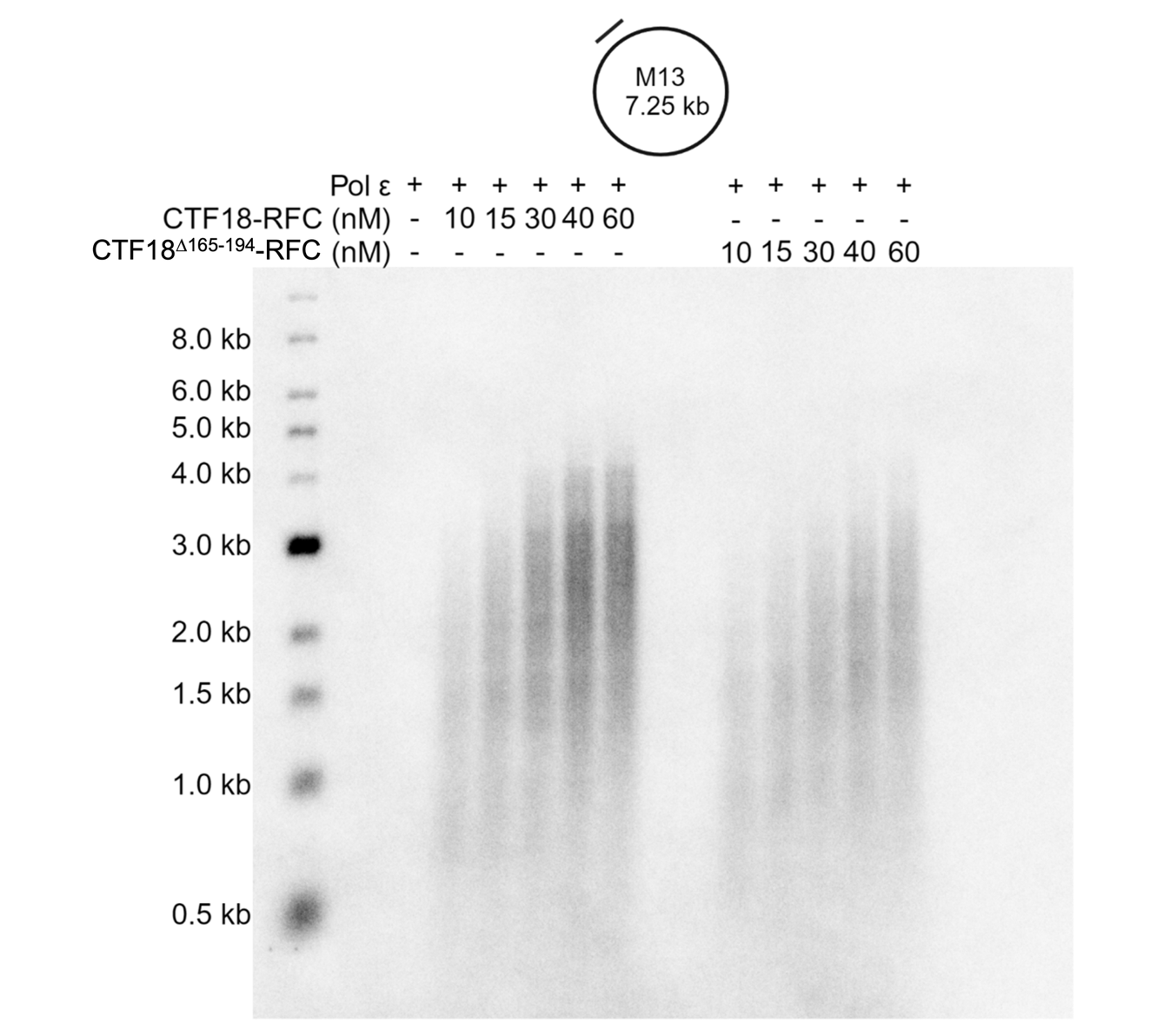

**S15 Fig.** *Primer extension assays with Pol ε and CTF18 WT and CTF18 ^Δ165-194^-RFC.* CTF18-RFC or *CTF18 ^Δ165-194^ -RFC* concentration titration reactions on M13mp18 single-strand DNA. Reactions were running for 10 mins at 37°C.

|  | CTF18-PCNA  with ATP | CTF18-PCNA  with ATP and Mg^2+^ |
| --- | --- | --- |
| **Data collection and processing** |  |  |
| Magnification | 130,000 | 130,000 |
| Voltage (kV) | 300 | 300 |
| Electron exposure (e–/Å^2^) | 43 | 46 |
| Defocus range (μm) | -2.7 to -1.5 | -2.7 to -1.5 |
| Pixel size (Å) | 0.93 | 0.93 |
| Symmetry imposed | C1 | C1 |
| Initial particle images (no.) | 2,043,120 | 3,216,167 |
| Final particle images (no.) | 527,527 | 602,591 |
| Map resolution (Å)  FSC threshold | 2.9  0.143 | 3.3  0.143 |
| Map resolution range (Å) | 2.8-3.8 | 3.1-4.3 |
| **Refinement** |  |  |
| Initial model used (PDB code) | 6VVO | 6VVO |
| Model resolution (Å)  FSC threshold | 3.0  0.5 | 3.5  0.5 |
| Map sharpening *B* factor (Å^2^) | -67.879 | -138.637 |
| Model composition  Non-hydrogen atoms  Protein residues  Ligands | 19777  2503  4 | 19779  2503  6 |
| *B* factors (Å^2^)  Protein  Ligand | 68.48  80.11 | 169.79  167.64 |
| R.m.s. deviations  Bond lengths (Å)  Bond angles (°) | 0.004  0.612 | 0.003  0.547 |
| Validation  MolProbity score  Clashscore  Poor rotamers (%) | 2.26  11.75  3.47 | 2.00  9.39  2.19 |
| Ramachandran plot  Favored (%)  Allowed (%)  Disallowed (%) | 96.14  3.82  0.04 | 96.38  3.49  0.12 |

**S1 Table.** Cryo-EM data collection, model refinement, and validation statistics.

**Supplementary Video 1.** *3DVA analysis of the human CTF18-RFC–PCNA complex in the presence of ATP.*
The video shows morphing transitions across the first three principal components obtained by 3D variability analysis (3DVA). Three volumes per component are morphed, illustrating continuous conformational flexibility within the complex. The dominant motion is localized to the AAA⁺ domain of the CTF18 subunit, consistent with reduced local resolution in this region and supporting a model in which this domain is intrinsically flexible within the assembled loader.

**References**

1.	Hedglin M, Perumal SK, Hu Z, Benkovic S. Stepwise assembly of the human replicative polymerase holoenzyme. Elife. 2013;2:e00278.

2.	Hedglin M, Benkovic SJ. Replication Protein A Prohibits Diffusion of the PCNA Sliding Clamp along Single-Stranded DNA. Biochemistry. 2017;56(13):1824-35.

3.	Hedglin M, Aitha M, Benkovic SJ. Monitoring the Retention of Human Proliferating Cell Nuclear Antigen at Primer/Template Junctions by Proteins That Bind Single-Stranded DNA. Biochemistry. 2017;56(27):3415-21.

4.	Kim C, Paulus BF, Wold MS. Interactions of human replication protein A with oligonucleotides. Biochemistry. 1994;33(47):14197-206.

5.	Kim C, Snyder RO, Wold MS. Binding properties of replication protein A from human and yeast cells. Mol Cell Biol. 1992;12(7):3050-9.

6.	Kim C, Wold MS. Recombinant human replication protein A binds to polynucleotides with low cooperativity. Biochemistry. 1995;34(6):2058-64.
